# Immune gene co-expression networks are constrained by a non-linear topology–spectral relationship

**DOI:** 10.64898/2025.12.25.696503

**Authors:** Roberto Navarro Quiroz, Eloina Zárate Peñate, Lisandro Pacheco Lugo, Yirys Díaz Olmos, Leonardo Pacheco Londoño, Antonio Acosta Hoyos, Elkin Navarro Quiroz

## Abstract

Inter-individual heterogeneity poses a fundamental challenge to systems-level characterization of immune states. When examined in isolation, network metrics exhibit extensive distributional overlap between conditions, providing little discriminatory power at the donor level. We hypothesized that immune states differ not in isolated properties but in the geometric constraints governing their joint distribution. We constructed donor-level gene co-expression networks from peripheral blood mononuclear cells across a Discovery cohort (CLARITY/SLE; n∼Healthy∼ = 74, n∼Disease∼ = 86) and an independent Validation cohort (STEPHENSON/COVID-19), totaling 259 unique donors. Network modularity (Q) and leading eigenvalue (λ∼max∼) distributions overlapped substantially between conditions (Q: Wilcoxon p = 0.913, Cliff’s δ = 0.01, 95% CI [−0.17, 0.19]; λ∼max∼: p = 0.108, δ = 0.15, 95% CI [−0.04, 0.34]). However, mapping donors into a joint modularity–spectral energy space revealed a non-monotonic organizational constraint: networks with high modularity exhibit constrained spectral energy, while low-modularity networks permit dominant collective modes. Critically, configurations combining high modularity with high spectral leakage are empirically absent—defining an **empirically unoccupied (’forbidden’) zone** in the state space. The two cohorts occupy opposite arms of this U-shaped crossover (Discovery: ρ = −0.39; Validation: ρ = +0.41), explaining their opposite correlation signs as sampling different regions of a single underlying constraint. This geometry persists across network densities (5–10%), sampling depths (500–800 cells), and within major cell lineages (CD4+ T cells, Monocytes). These findings characterize immune network organization not by univariate biomarkers, but by geometric boundaries that constrain permissible configurations.

**Significance Statement:** Immune responses vary substantially across individuals, frustrating efforts to identify universal organizational principles. We analyzed gene co-expression networks from peripheral blood mononuclear cells across 259 donors in two independent cohorts and found that neither network modularity nor spectral dominance alone distinguishes healthy from disease states—distributions overlap extensively. However, plotting these properties jointly reveals that observed immune networks are confined to a restricted region of the modularity–spectral energy space, while configurations combining high modularity with high spectral energy are empirically absent. This geometric constraint persists across cohorts, sampling depths, and cell lineages, suggesting a **robust structural trade-off** between compartmentalization and collective coordination in immune network architecture.

## Introduction

The immune system operates as a distributed network of interacting cells and molecules, where coordinated gene expression programs underlie both homeostatic function and pathological responses. Yet individual variation—arising from genetic background, environmental exposures, disease history, and stochastic developmental processes—generates substantial heterogeneity in any population-level measurement (1, 2). This heterogeneity has frustrated efforts to identify universal organizational principles of immune dysfunction, as metrics that appear to discriminate at the group level often exhibit extensive overlap when examined at the individual level (3, 4).

Reductionist approaches that focus on single biomarkers or isolated network properties face inherent limitations when confronting this heterogeneity. Network modularity—quantifying the extent to which genes cluster into densely connected communities—has been widely adopted as a summary statistic for co-expression architecture (5–7). Similarly, spectral analysis of correlation matrices, particularly the examination of dominant eigenvalues, provides information about collective expression modes that span network modules (8–10). Random matrix theory offers analytical expectations for eigenvalue distributions in noise-dominated systems, against which empirical spectra can be compared to distinguish signal from stochastic fluctuations (11–13). However, these properties are typically analyzed in isolation, and their failure to robustly discriminate between conditions is often attributed to noise rather than examined as a potentially informative pattern.

A complex systems perspective suggests an alternative interpretation. Even when two metrics individually overlap between groups, their joint distribution may reveal regime-specific constraints invisible to univariate analysis (14–16). The relationship between network topology and collective dynamics—how modular architecture constrains or permits coordinated expression patterns—represents a coupling that could characterize organizational states more robustly than either property alone. Such constraints, if they exist, would define not merely correlations but geometric boundaries: regions of the state space that are biologically permissible versus those that are structurally forbidden.

This perspective motivates a shift from seeking marginal differences toward examining the geometry of permissible states. Complex adaptive systems—from neural networks to ecosystems—often exhibit trade-offs between compartmentalization and global coordination (17–19). Modular organization enables specialized, localized processing but limits system-wide synchronization; conversely, reduced modularity permits collective modes but sacrifices compartmental integrity. If immune networks obey similar organizational principles, we might expect their configurations to cluster along characteristic trajectories rather than filling the modularity–spectral space uniformly.

We do not aim to identify biomarkers, predict clinical outcomes, or attribute mechanisms to specific molecular pathways. Rather, we test a focused hypothesis: that immune states are constrained by a non-monotonic geometric relationship between network topology and spectral structure. We construct donor-level co-expression networks from two independent cohorts, map their configurations into a joint modularity–spectral energy space, and examine whether this space exhibits structural boundaries—regions that are occupied versus those that remain empirically absent. The donor serves as the unit of inference throughout. Our goal is to characterize organizational constraints, with full acknowledgment that observational data cannot establish causality and that replication across cohorts tests robustness but not universality.

## Results

### Univariate Network Properties Fail to Discriminate Immune States at the Donor Level

We first assessed whether immune states could be distinguished by their global network topology or spectral properties when examined independently. Conceptual modeling proposes that immune dysfunction involves a transition from a modular, compartmentalized architecture toward a globally coupled organization where dominant collective modes emerge (Fig. 1A). **This schematic is illustrative only and is not derived from empirical data; it serves solely to frame the organizational hypothesis under test.**

**Figure 1.**
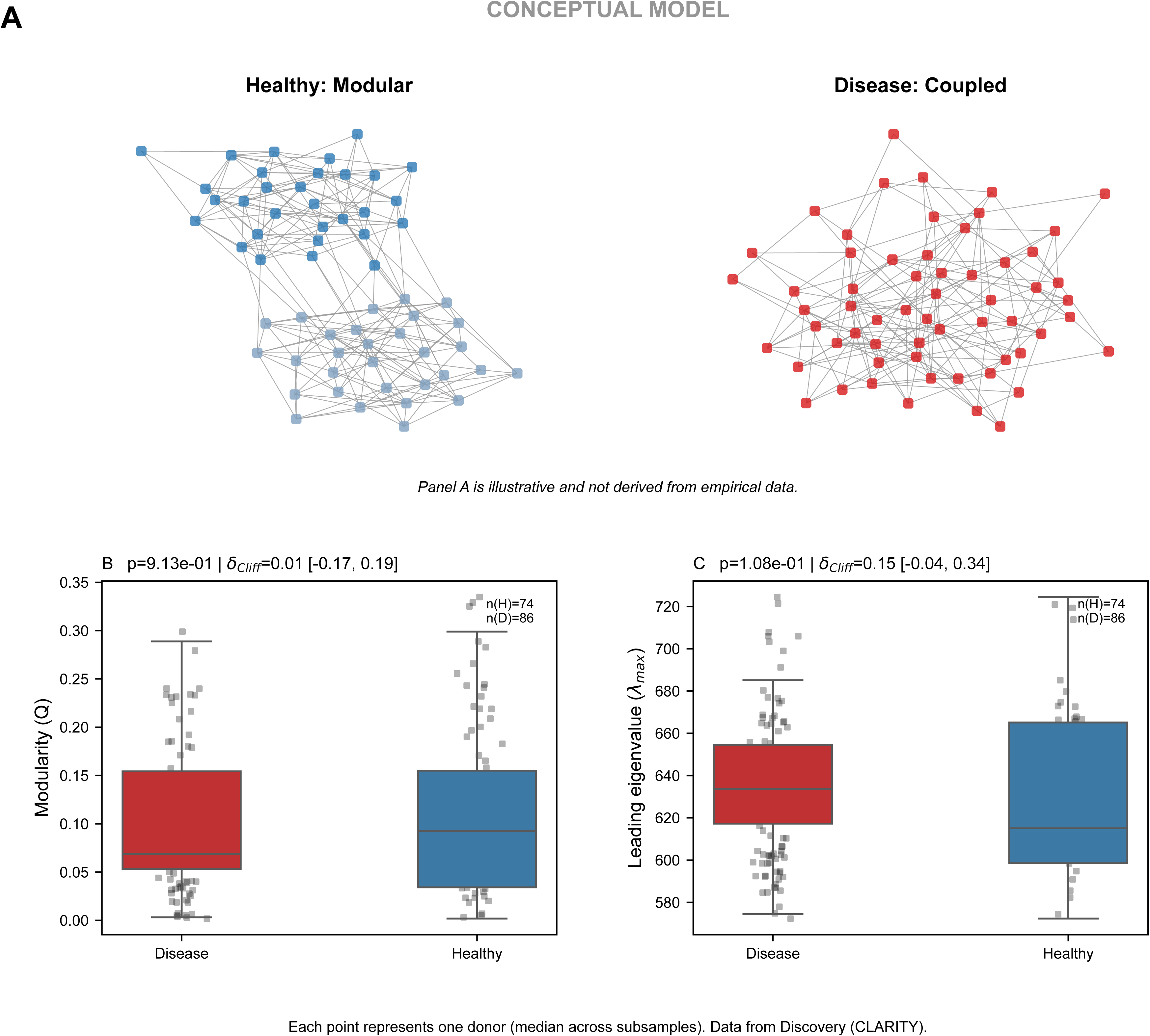
Univariate network properties fail to discriminate immune states at the donor level. (A) Conceptual model (ILLUSTRATIVE ONLY). Schematic depicting hypothesized transition from modular to globally coupled network architecture. This panel is not derived from empirical data; it serves solely to frame the organizational hypothesis under test. (B) Distribution of network modularity (Q) across disease (n = 86, red) and healthy (n = 74, blue) donors in the Discovery cohort (CLARITY). Each point represents one donor (median across subsamples). Wilcoxon rank-sum test: p = 0.913; Cliff’s δ = 0.01 (95% CI: [−0.17, 0.19]). (C) Distribution of leading eigenvalue (λ∼max∼) across conditions. Wilcoxon test: p = 0.108; Cliff’s δ = 0.15 (95% CI: [−0.04, 0.34]). Confidence intervals span zero in both cases, indicating no reliable discrimination.

Quantitative analysis of the Discovery cohort (CLARITY; n∼Healthy∼ = 74, n∼Disease∼ = 86 donors) revealed high inter-donor heterogeneity in both network modularity (Q) and the leading eigenvalue (λ∼max∼). Modularity distributions overlapped substantially between healthy and disease donors (Fig. 1B). A two-sided Wilcoxon rank-sum test yielded p = 0.913, indicating no detectable distributional difference. The effect size was negligible: Cliff’s δ = 0.01 (95% CI: [−0.17, 0.19]). The confidence interval spans zero symmetrically, providing no evidence that modularity differs between conditions at the population level.

Similarly, the leading eigenvalue showed weak distributional differences with no reliable separation (Fig. 1C). Wilcoxon rank-sum test: p = 0.108; Cliff’s δ = 0.15 (95% CI: [−0.04, 0.34]). While the point estimate suggests a modest effect favoring higher λ∼max∼ in disease, the confidence interval spans zero, precluding reliable inference of a condition difference.

Each point in Fig. 1B–C represents one donor, computed as the median value across subsampled instances (N = 500 and N = 800 cells per instance) to ensure robustness against technical variation. The substantial overlap reflects genuine inter-individual heterogeneity rather than measurement noise. These results establish that neither modularity nor spectral dominance alone serves as a discriminating feature when examined independently—justifying the need to examine their joint distribution.

### The Geometry of Permissible States: A Non-Monotonic Organizational Constraint

The principal finding emerged when mapping donors into a joint topology–spectral state space (Fig. 2). For each donor, we computed spectral energy outside an empirical null (Σλ∼out∼), defined as the sum of all eigenvalues exceeding the 95th percentile of a permutation-derived threshold. This permutation null was constructed by independently shuffling gene labels across cells for each donor (1,000 permutations), preserving global variance while destroying co-expression structure. The quantity Σλ∼out∼ thus captures variance attributable to collective modes beyond random expectation.

**Figure 2.**
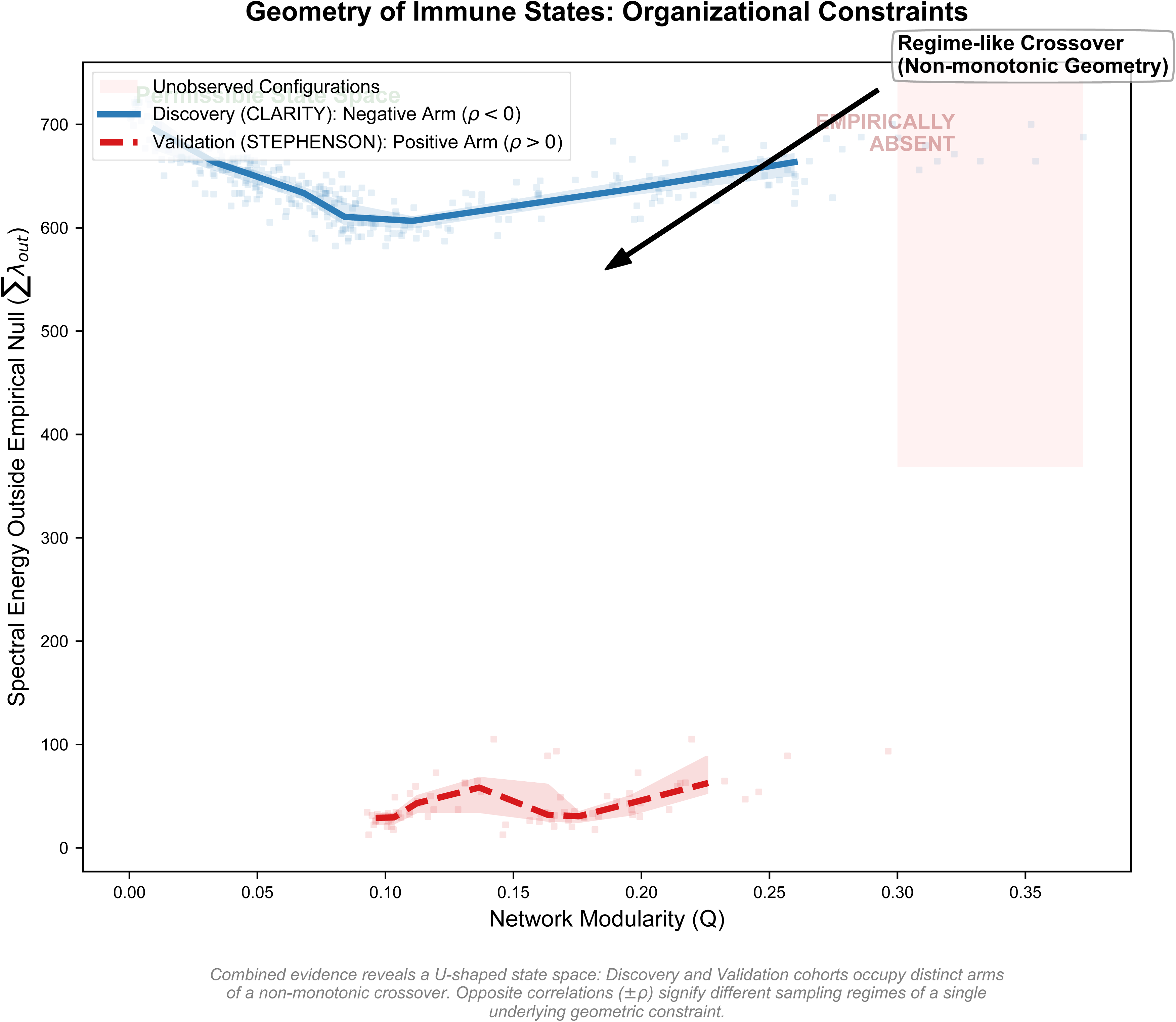
The geometry of permissible states: a non-monotonic organizational constraint. Joint distribution of spectral energy outside empirical null (Σλ∼out∼) versus network modularity (Q) for Discovery (CLARITY; solid blue line, blue points) and Validation (STEPHENSON; dashed red line, pink points) cohorts. Lines connect binned medians (n = 8 equal-count intervals); shaded regions indicate bootstrap 95% confidence intervals (2,000 iterations). Discovery: Spearman ρ = −0.39; Validation: ρ = +0.41. The opposite correlation signs reflect sampling of different arms of a U-shaped crossover, not inconsistency. Pink-shaded region labeled "EMPIRICALLY ABSENT" indicates configurations combining high modularity with high spectral energy that are not observed across the 259-donor meta-cohort—the geometric "forbidden zone." Arrow indicates regime-like crossover zone. Σλ∼out∼ defined as sum of eigenvalues exceeding 95th percentile of donor-specific permutation null (1,000 permutations per donor).

Plotting Σλ∼out∼ against modularity Q revealed a striking geometric pattern. Observed immune networks do not fill the Q–Σλ∼out∼ space uniformly. Instead, they cluster along a U-shaped trajectory that defines the **permissible state space** of immune network organization. At low modularity (Q < 0.08), networks exhibit elevated spectral energy outside the null, indicating dominant collective modes spanning the network. As modularity increases toward intermediate values (Q ≈ 0.10–0.15), spectral energy decreases, reaching a local minimum—the crossover zone where compartmentalization and collective coordination trade off most sharply. Beyond this zone, spectral energy stabilizes or shows modest recovery.

Critically, configurations combining high modularity (Q > 0.25) with high spectral energy are **empirically absent** (Fig. 2, pink-shaded region labeled "EMPIRICALLY ABSENT"). **This empirically unoccupied region represents network configurations that—while geometrically possible—are not observed in our meta-cohort of 259 donors under the immune contexts and sampling conditions examined.** The absence of such configurations suggests a structural constraint: high topological compartmentalization effectively suppresses the emergence of system-wide collective modes.

The two cohorts occupy opposite arms of this U-shaped constraint. The Discovery cohort (CLARITY) samples primarily the negative arm, where modularity and spectral energy are inversely related (Spearman ρ = −0.39, permutation p < 0.001). The Validation cohort (STEPHENSON) samples primarily the positive arm, where modest increases in modularity accompany increases in spectral energy (ρ = +0.41, permutation p < 0.001).

**The opposite signs of these correlations do not indicate inconsistency; they reveal the non-monotonic geometry.** A linear correlation coefficient is an inadequate descriptor of a U-shaped relationship. The cohorts differ in their sampling position along the modularity axis and in absolute scale of Σλ∼out∼ (Discovery: 580–720; Validation: 20–100)—differences likely arising from technical variation in network construction, cell sampling, and disease composition. What is preserved across cohorts is the geometric shape: both trace constrained trajectories rather than filling the space, and both respect the same forbidden zone.

To visualize this pattern while accounting for heteroscedasticity, we computed binned medians using eight equal-count intervals across the Q range, with bootstrap 95% confidence intervals (2,000 iterations). The coupling relationship was stable across different binning counts (6–10 bins) and preserved under leave-one-donor-out robustness analysis, confirming that no single donor drives the observed pattern.

### The Organizational Constraint Persists Across Technical Parameters

A critical concern is whether the observed geometry reflects genuine biological organization or artifacts of specific analytical choices. We addressed this through systematic sensitivity analysis.

### Network density threshold

The U-shaped constraint persists when networks are constructed at both 5% density (main analysis) and 10% density (Fig. S1A–B). Although absolute modularity values shift predictably with density, the non-monotonic shape and the empirically absent region remain invariant. This demonstrates that the constraint is not an artifact of a particular threshold choice.

### Sampling depth

The constraint remains stable when computed from subsamples of 500 cells versus 800 cells per donor instance (Fig. S2A–B). Despite expected changes in absolute magnitudes due to varying information density, the functional form of the relationship persists. This confirms that the geometry reflects genuine collective organization rather than sampling noise.

### Cell lineage composition

To address whether the constraint merely reflects varying proportions of cell types across donors, we repeated the analysis within computationally purified populations of abundant cell lineages: CD4+ T cells (Fig. S3A) and Monocytes (Fig. S3B). The non-monotonic constraint replicates within these lineage-restricted networks, albeit with lineage-specific shifts in absolute position. This demonstrates that the organizational constraint is an intrinsic property of co-expression architecture, not solely driven by compositional variation.

#### Spectral Fingerprinting Validates the Σλ∼out∼ Metric

To further characterize the spectral structure underlying the geometric constraint, we performed spectral fingerprinting by comparing observed eigenvalue distributions to the permutation-derived empirical null (Fig. 3). The 95th percentile of the null distribution served as the primary threshold for classifying eigenvalues as non-random; the theoretical Marčenko–Pastur edge was provided as secondary visual reference only and was not used for inference (20).

**Figure 3.**
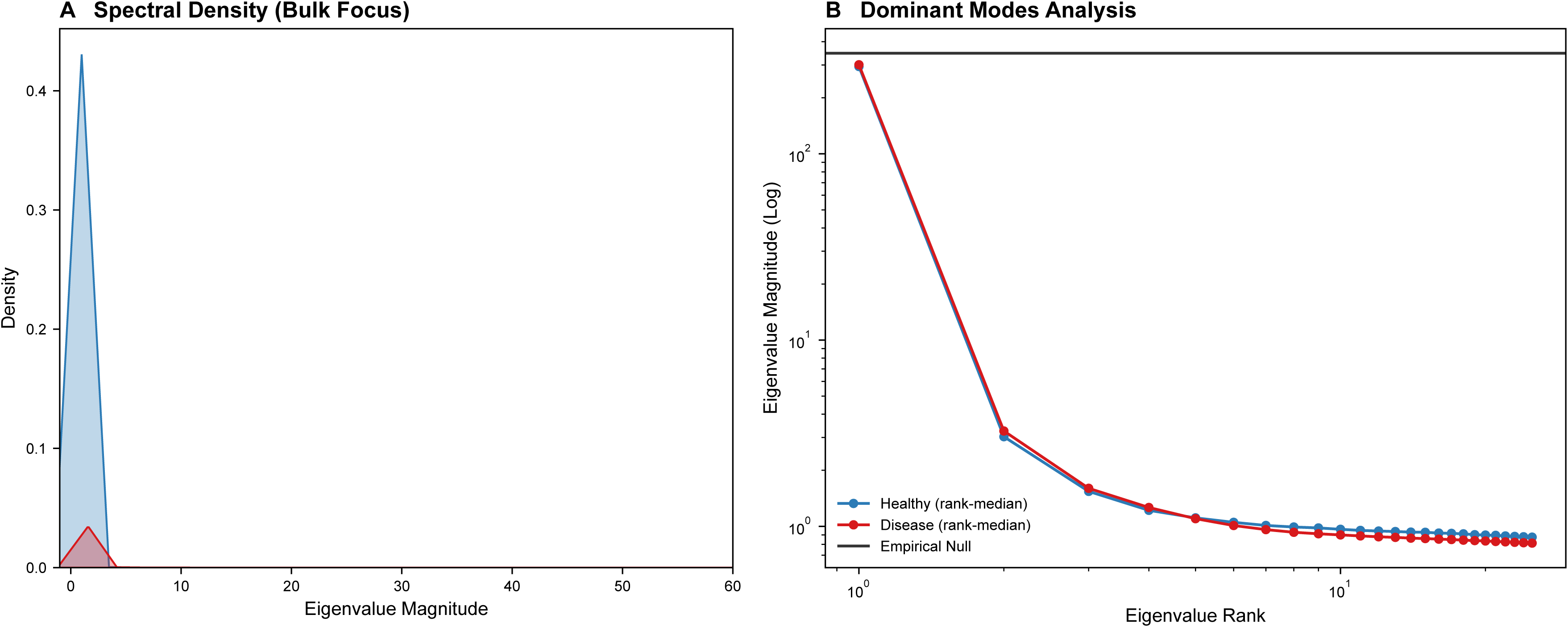
Spectral fingerprinting validates collective modes exceeding empirical null. (A) Bulk spectral density distribution comparing healthy (blue) and disease (red) networks. Most eigenvalues cluster near zero; disease shows slightly broader distribution extending to higher magnitudes. (B) Dominant modes analysis on log-log axes. Points show rank-median eigenvalue magnitudes across donors for healthy (blue) and disease (red). Black line indicates empirical null threshold (95th percentile of permutation distribution). Both conditions exhibit eigenvalues substantially exceeding null expectation, confirming non-random collective modes. The Marčenko–Pastur theoretical edge serves only as secondary visual reference; all inferences use the permutation-derived empirical null.

The bulk spectral density (Fig. 3A) showed that most eigenvalues cluster near zero in both conditions, reflecting variance attributable to pairwise correlations without collective organization. Disease networks exhibited a slightly broader distribution, with more density extending toward higher eigenvalue magnitudes, though bulk structures substantially overlap.

Dominant modes analysis (Fig. 3B) plotted eigenvalue magnitude against rank on logarithmic axes. Both healthy and disease networks showed rank-median eigenvalue profiles substantially exceeding the empirical null threshold (black line), confirming that observed networks possess collective modes unexplained by random correlation structure. Disease networks showed modestly elevated eigenvalues at the highest ranks, indicating a more pronounced hierarchy of dominant modes—consistent with the elevated Σλ∼out∼ values observed in low-modularity configurations.

The persistence of eigenvalues above the empirical null across different subsampling sizes confirms that this spectral structure is robust to technical variation. Both conditions exhibit non-random spectral organization; the distinction lies not in presence versus absence of collective modes, but in how their magnitude relates to topological modularity—the geometric coupling captured in Fig. 2.

## Discussion

### The Geometry of Organizational Constraints

We report a non-monotonic geometric constraint governing immune gene co-expression networks. This constraint emerges despite the absence of detectable group differences when topology or spectral properties are examined independently. The key finding is not a correlation but a geometry: immune networks occupy a restricted region of the modularity–spectral energy space, while configurations combining high compartmentalization with high collective dominance are empirically absent.

The physical interpretation is intuitive within complex systems theory (21–23). Networks with low modularity lack strong topological partitions, permitting collective modes to span the entire system; such configurations exhibit high spectral energy outside the random bulk. As modularity increases, community boundaries emerge that constrain global coordination, reducing variance captured by system-wide modes. **The empirically unoccupied region corresponding to high modularity with high spectral energy represents an organizational configuration that is not realized in the immune systems examined here.** We emphasize that ’forbidden’ is used here in a descriptive, empirical sense to denote regions of the topology–spectral space that remain unoccupied in the datasets analyzed, not as a claim of mathematical impossibility or universal biological law.

This interpretation frames the constraint as a trade-off between compartmentalization and synchronization, analogous to constraints observed in other complex adaptive systems.

Neural networks exhibit similar trade-offs between modular processing and global integration (17, 24, 25). Ecological food webs balance compartmentalization—which provides robustness against local perturbations—with connectedness that permits systemic responses (18, 26).

The immune system, operating as a distributed information-processing network, may obey related architectural principles.

### Resolving the Paradox of Signs Through Geometric Interpretation

The opposite signs of Spearman correlations between cohorts (Discovery: ρ = −0.39; Validation: ρ = +0.41) initially appear contradictory. Our geometric interpretation resolves this paradox: both cohorts sample the same underlying U-shaped constraint but occupy different arms of the crossover.

The sign of a linear correlation coefficient depends on which portion of a non-monotonic curve is predominantly sampled. If sampling concentrates on the left arm (low Q, decreasing Σλ∼out∼ with increasing Q), ρ will be negative. If sampling concentrates on the right arm (higher Q, increasing Σλ∼out∼ with increasing Q), ρ will be positive. This is precisely what we observe.

The scale differences between cohorts (Discovery: Σλ∼out∼ ≈ 580–720; Validation: Σλ∼out∼ ≈ 20–100) likely reflect technical variation in network construction parameters, gene selection, and disease heterogeneity. What is preserved is the shape invariance: the non-monotonic functional form and the empirically absent region. Cross-cohort quantitative comparison requires harmonization that was not attempted here; our claim is limited to shape replication, which provides stronger evidence for a generalizable organizational principle than quantitative identity would.

### Relation to Prior Work

Network modularity has been extensively studied in biological systems, from protein interaction networks to brain connectomes (5, 6, 27–29). Spectral analysis of correlation matrices has complementary origins in random matrix theory, initially developed for nuclear physics and subsequently applied to financial, neural, and genomic systems (11–13, 30, 31). Our contribution integrates these traditionally separate frameworks by examining their joint distribution and, crucially, by identifying geometric constraints rather than merely reporting correlations.

Prior studies of immune co-expression networks have documented condition-associated changes in modularity or hub structure (32–34), but typically report univariate comparisons that—as we demonstrate—exhibit extensive overlap at the donor level. By shifting focus to geometric constraints, we reveal structure invisible to marginal analyses. The absence of certain configurations (the forbidden zone) is itself an empirical finding that univariate approaches cannot detect.

The application of permutation-based empirical nulls, rather than theoretical RMT distributions, addresses a practical limitation: biological correlation matrices violate independence assumptions underlying analytical results like the Marčenko–Pastur law (20). Permutation nulls provide data-driven thresholds appropriate for each donor’s specific covariance structure, improving specificity at the cost of computational expense.

### Biological Interpretation: Speculative but Constrained

From an immunological perspective, the coupling between modularity and spectral dominance admits biological interpretation, though we emphasize this remains speculative without experimental validation. Modular co-expression architecture may reflect functional compartmentalization: distinct immune cell subsets or transcriptional programs operating semi-independently (35, 36). High modularity could indicate that lymphocyte, myeloid, and other compartments maintain distinct expression programs with limited cross-talk.

Conversely, reduced modularity with elevated spectral energy may indicate increased global coordination—potentially reflecting inflammatory states where systemic signals (cytokines, interferons) drive correlated responses across cell types (37, 38). The dominant spectral modes exceeding random expectation could capture this system-wide coordination. Such interpretation is consistent with concepts of cytokine storm or systemic inflammatory response, though we caution against over-interpretation: bulk transcriptomic data cannot resolve whether observed correlations reflect within-cell regulatory coupling, compositional shifts, or technical confounding.

The forbidden zone—high modularity with high spectral energy—may represent an organizational impossibility in immune systems. If compartmentalization requires insulation between modules, it structurally limits the emergence of modes that span modules.

Conversely, if systemic coordination requires breakdown of modular boundaries, high modularity and high collective dominance cannot coexist. This interpretation, while speculative, generates falsifiable predictions for future experimental work.

### Limitations

Several limitations constrain interpretation and should be transparently acknowledged:

### Observational design

This study is purely observational. We cannot establish whether topological changes precede, follow, or co-occur with spectral changes, nor can we determine causal relationships between network architecture and immune function. The constraint describes correlation structure, not mechanism.

### Bulk transcriptomics aggregates cell types

Observed modularity and spectral properties aggregate across all cell types present in peripheral blood. Changes in cell-type composition between conditions could contribute to apparent network reorganization. While our lineage-specific analysis (Fig. S3) demonstrates that the constraint persists within CD4+ T cells and Monocytes, this does not fully exclude compositional confounding, and single-cell deconvolution approaches would strengthen the analysis (39, 40).

### Technical parameter sensitivity

Network construction involves choices—correlation metric, thresholding strategy, gene selection—that affect absolute values. Our sensitivity analyses demonstrate shape invariance across parameters, but we cannot exhaustively test all analytical variants. The scale differences between cohorts highlight this sensitivity.

### Heterogeneity remains substantial

Even with the geometric constraint, individual donors exhibit wide scatter around the binned medians (Fig. 2). The constraint characterizes population-level organizational boundaries, not individual-level classification. Application to individual prediction would require demonstration of discriminative power not attempted here.

### Unknown generalizability

Two cohorts from related contexts (PBMC in inflammatory conditions) cannot establish universality. Whether the constraint holds in other tissues, species, or disease contexts is unknown. Extension to additional cohorts sampling the full modularity range with harmonized parameters is required.

### Broader Implications

The geometry of permissible states reported here may exemplify principles extending beyond immune networks. Complex adaptive systems across domains exhibit trade-offs between modularity and integration, between specialized processing and global coherence. The finding that certain configurations are empirically absent—not merely rare, but structurally forbidden—**suggests constraints on how such systems may be organized under comparable architectural and dynamical conditions.**

From a methodological standpoint, our findings illustrate the value of examining joint distributions rather than marginal statistics, and of identifying geometric constraints rather than merely correlations. The null results for univariate comparisons (Fig. 1B–C) could be dismissed as "no difference"—yet the geometric analysis (Fig. 2) reveals structure these comparisons miss. This argues for routine examination of higher-order relationships in systems biology, particularly when univariate heterogeneity obscures simple group differences.

We emphasize that these findings do not enable clinical application. The constraint characterizes organizational principles at the population level; implications for diagnosis, prognosis, or therapy remain unexplored. Our contribution is conceptual: demonstrating that immune network organization obeys geometric boundaries that define permissible configurations and exclude others, offering a falsifiable framework for understanding complex biological systems.

## Materials and Methods

### Data Sources and Cohorts

Two independent single-cell RNA sequencing datasets of peripheral blood mononuclear cells (PBMCs) were analyzed:

### Discovery cohort (CLARITY/SLE)

n∼Healthy∼ = 74 donors, n∼Disease∼ = 86 donors with systemic lupus erythematosus. Data from Perez et al. (41), available through GEO under accession GSE174188. Quality control, normalization, and cell-type annotation followed original study protocols.

### Validation cohort (STEPHENSON/COVID-19)

Independent PBMC dataset from Stephenson et al. (42), available through ArrayExpress under accession E-MTAB-10026, comprising donors with COVID-19 and healthy controls.

The combined meta-cohort comprised 259 unique donors.

### Unit of Analysis

The individual donor served as the statistical unit throughout. All comparative statistics (Wilcoxon tests, effect sizes, correlations) were computed on donor-level aggregated values, not cells. This ensures that biological conclusions reflect inter-individual variation rather than pseudo-replication from correlated cells within donors (43).

### Subsampling Strategy

To ensure robustness against technical noise and variable cell counts, each donor was subsampled multiple times (N = 2 instances: 500 and 800 cells per instance). Network metrics were computed for each subsample; the median across subsamples served as the donor-level estimate. This approach mitigates sensitivity to cell count variation and provides stability against sampling fluctuation. Donors with fewer than the minimum required cells (500) were excluded.

### Network Construction

For each donor-subsample, a gene × gene correlation matrix R was computed using Pearson correlation coefficients between highly variable gene pairs. To standardize network density and reduce noise, a density-based threshold retained only the top 5% of correlations (by absolute magnitude) as edges. The resulting adjacency matrix was unweighted (binary). This threshold was held constant across all donors, cohorts, and conditions to prevent density-driven confounding. Robustness was verified at 10% density (Fig. S1).

### Modularity (Q)

Network modularity was computed using the Louvain algorithm (44), which maximizes: Q = (1/2m) Σ∼ij∼ [A∼ij∼ − k∼i∼k∼j∼/2m] δ(c∼i∼, c∼j∼) where A is the adjacency matrix, k∼i∼ is the degree of node i, m is total edges, and δ(c∼i∼, c∼j∼) = 1 if nodes i and j belong to the same community. Resolution parameter γ = 1.0 was fixed throughout; no per-condition tuning was performed to ensure falsifiability.

### Spectral Analysis

Eigenvalue decomposition of the correlation matrix R yielded eigenvalues λ∼1∼ ≥ λ∼2∼ ≥ … ≥ λ∼n∼. Two metrics were extracted:

### Leading eigenvalue (λ∼max∼)

The largest eigenvalue, representing the dominant collective mode’s magnitude.

### Spectral energy outside empirical null (Σλ∼out∼)

Defined as Σ∼i∼ λ∼i∼ · I(λ∼i∼ > τ), where τ is the 95th percentile of the permutation-derived null eigenvalue distribution and I(·) is the indicator function. This metric quantifies total variance attributable to collective modes exceeding random expectation.

### Empirical Null Model

For each donor, a permutation null was constructed by shuffling gene labels across cells independently (1,000 permutations), preserving row and column marginals while destroying correlation structure. The eigenvalue distribution of permuted matrices defined the null. The 95th percentile of this distribution served as the threshold τ for classifying eigenvalues as non-random. The theoretical Marčenko–Pastur edge (20) was computed for visualization but not used for inference.

### Statistical Analysis

#### Group comparisons

Distributions of Q and λ∼max∼ between conditions were compared using two-sided Wilcoxon rank-sum tests. Effect sizes were quantified using Cliff’s δ (45), a non-parametric measure ranging from −1 to +1, with 95% confidence intervals computed via 10,000 bootstrap resamples.

### Coupling analysis

The relationship between Q and Σλ∼out∼ was assessed using Spearman rank correlation with permutation-based p-values (5,000 permutations). To visualize non-linear structure, donors were binned into n = 8 equal-count intervals by Q; bin medians and bootstrap 95% CIs (2,000 iterations) were computed.

### Robustness

Leave-one-donor-out analysis confirmed that no single donor drove the coupling pattern. Sensitivity analyses across density thresholds (5%, 10%) and sampling depths (500, 800 cells) verified shape invariance.

### Lineage-Specific Analysis

To control for compositional confounding, networks were reconstructed using only cells from specific major lineages (CD4+ T cells, Monocytes) where sufficient cell counts permitted (Fig. S3).

### Reproducibility

All stochastic processes (subsampling, permutations, bootstrapping) used fixed random seeds to ensure exact reproducibility. Analysis code will be made publicly available upon acceptance.

## Data Availability

All datasets analyzed in this study are publicly available. The SLE PBMC dataset from Perez et al. (41) is available through GEO under accession GSE174188. The COVID-19 PBMC dataset from Stephenson et al. (42) is available through ArrayExpress under accession E-MTAB-10026. All scripts for data processing, analysis, and figure generation will be made publicly available upon manuscript acceptance.

## Acknowledgments

The authors thank the Colombian Association of Immunology (ACOI) for sustained academic support, scientific discussion, and community building that contributed to the development of this work. This research was supported by Universidad Simón Bolívar and the Centro de Investigaciones en Ciencias de la Vida (CICV).

## Supplementary Information

**Figure S1.**
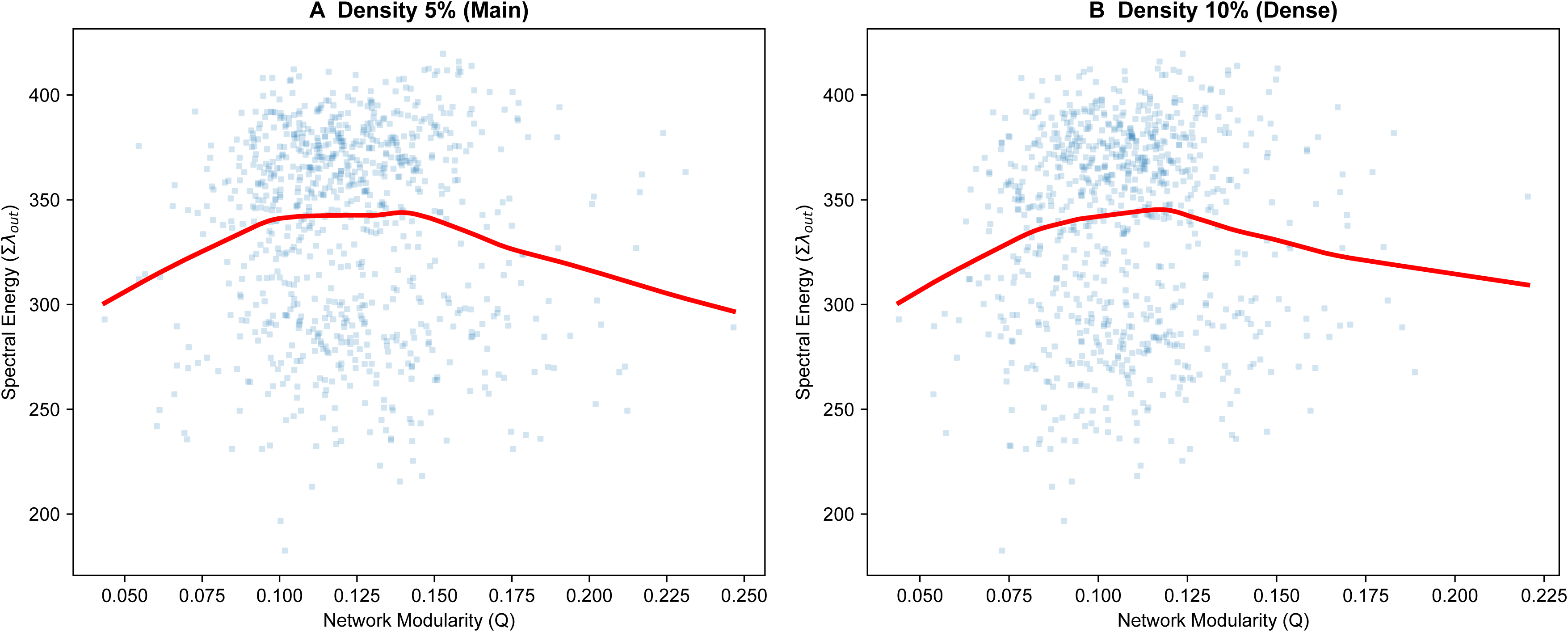
Robustness of the organizational constraint across network density thresholds. (A–B) The joint distribution of modularity (Q) and spectral energy (Σλ∼out∼) is shown for donor-level networks constructed at varying edge densities: (A) 5% (main text parameter) and 10% (dense coverage). The characteristic non-monotonic trajectory and regime crossover are preserved across densities. Although absolute modularity values shift predictably with density, the geometric shape of the constraint remains invariant, demonstrating that the phenomenon is not an artifact of a specific threshold choice.

**Figure S2.**
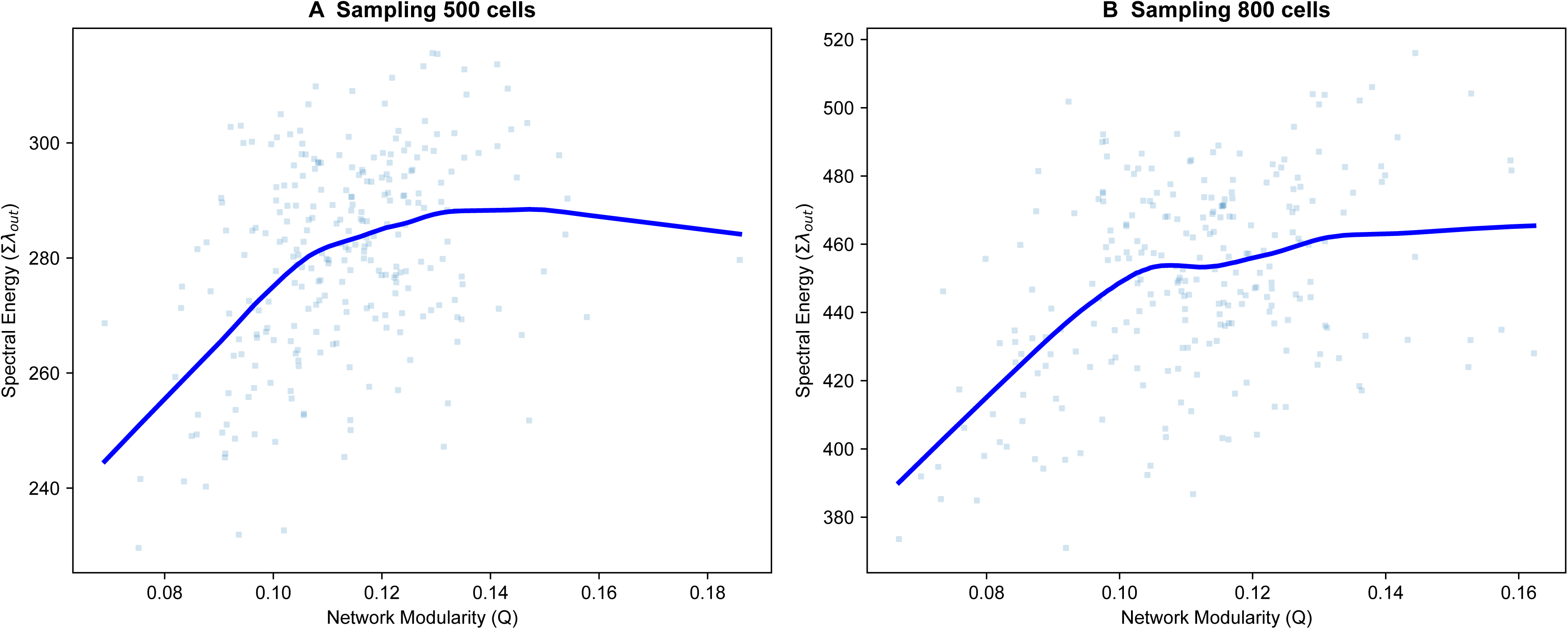
Stability of the organizational constraint against sampling depth. (A–B) The coupling analysis was repeated using different levels of subsampling depth: (A) 500 cells and (B) 800 cells per donor instance. Despite expected changes in the absolute magnitude of spectral energy and modularity due to varying density of information, the non-linear geometric form of the relationship persists. This confirms that the constraint is a robust property of the system’s collective mode structure, not a trivial consequence of the number of cells analyzed.

**Figure S3.**
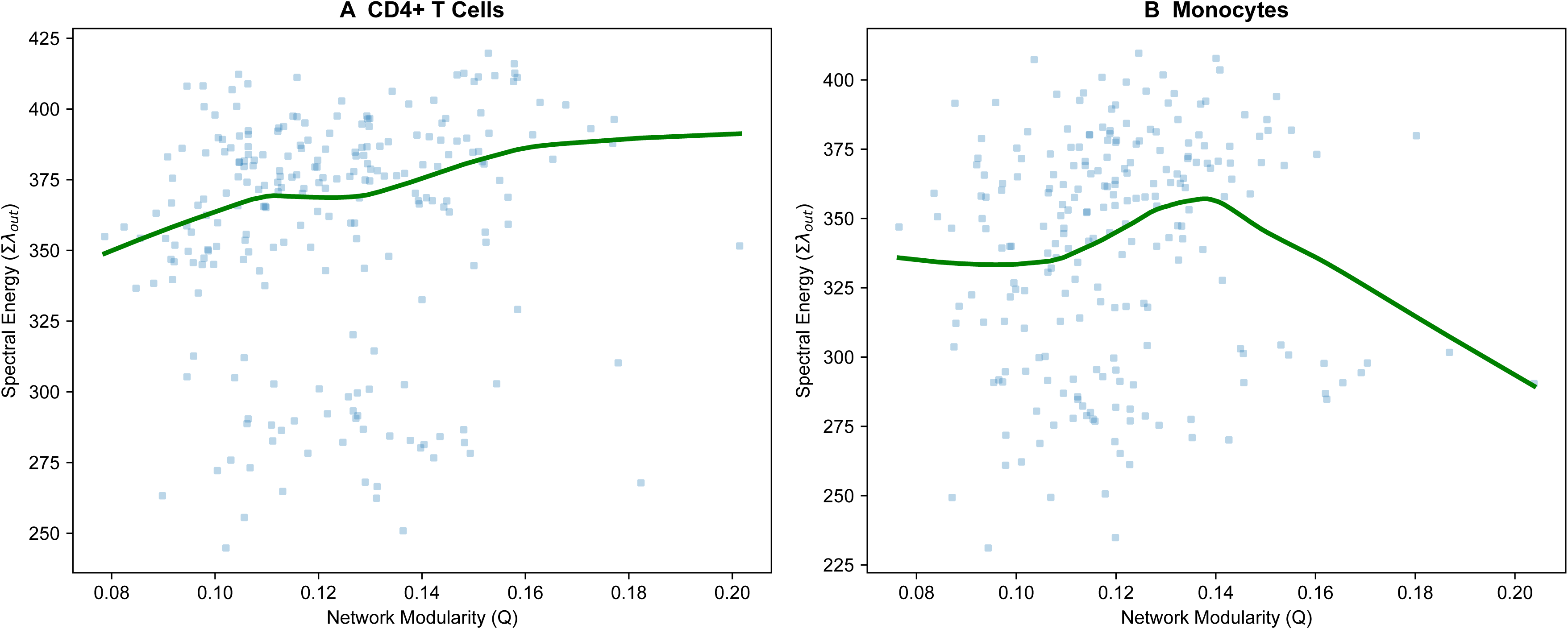
The organizational constraint persists within major cell lineages. (A–B) To control for the effect of inter-donor variation in cell-type frequencies (compositional bias), networks were reconstructed using only cells from specific major lineages: (A) CD4+ T cells and (B) Monocytes. The non-monotonic constraint replicates within these computationally purified populations, albeit with lineage-specific shifts in the absolute position within the state space. This result indicates that the observed constraint reflects genuine regulatory architecture rather than being driven solely by changes in the proportion of cell types.

